# Comparative genomics analysis reveals high levels of differential DNA transposition among primates

**DOI:** 10.1101/520387

**Authors:** Wanxiangfu Tang, Ping Liang

## Abstract

Mobile elements generated via DNA transposition constitute ∼50% of the primate genomes. As a result of past and ongoing activity, DNA transposition is responsible for generating inter- and intra-species genomic variations, and it plays important roles in shaping genome evolution and impacting gene function. While limited analysis of mobile elements has been performed in many primate genomes, a large-scale comparative genomic analysis examining the impact of DNA transposition on primate evolution is still missing.

Using a bioinformatics comparative genomics approach, we performed analysis of species-specific mobile elements (SS-MEs) in eight primate genomes, which include human, chimpanzee, gorilla, orangutan, green monkey, crab-eating macaque, rhesus monkey, and baboon. These species have good representations for the top two primate families, *Hominidae* (great apes) and the *Cercopithecidae* (old world monkeys), for which draft genome sequences are available.

Our analysis identified a total of 230,855 SS-MEs from the eight primate genomes, which collectively contribute to ∼82 Mbp genome sequences, ranging from 14 to 25 Mbp for individual genomes. Several new interesting observations were made based on these SS-MEs. First, the DNA transposition activity level reflected by the numbers of SS-MEs was shown to be drastically different across species with the highest (baboon genome) being more than 30 times higher than the lowest (crab-eating macaque genome). Second, the compositions of SS-MEs, as well as the top active ME subfamilies, also differ significantly across genomes. By the copy numbers of SS-MEs divided into major ME classes, SINE represents the dominant class in all genomes, but more so in the *Cercopithecidae* genomes than in the *Hominidae* genomes in general with the orangutan genome being the outliner of this trend by having LINE as the dominant class. While *AluY* represents the major SINE groups in the *Hominidae* genomes, *AluYRa1* is the dominant SINE in the *Cercopithecidae* genomes. For LINEs, each *Hominidae* genome seems to have a unique most active *L1* subfamily, but all *Cercopithecidae* genomes have L1RS2 as the most active LINEs. While genomes with a high number of SS-MEs all have one or more very active ME subfamilies, the crab-eating macaque genome, being the one with an extremely low level of DNA transposition, has no single ME class being very active, suggesting the existence of a genome-wide mechanism suppressing DNA transposition. Third, DNA transposons, despite being considered dead in primate genomes, were in fact shown to have a certain level of activity in all genomes examined with a total of ∼2,400 entries as SS-MEs. Among these SS-MEs, at least 23% locate to genic regions, including exons and regulatory elements, presenting significant potentials for their impact on gene function. Very interestingly, our data demonstrate that, among the eight primates included in this study, the human genome is shown to be the most actively evolving genome via DNA transposition as having the highest most recent activity of many ME subfamilies, notably the AluYa5/Yb8/Yb9, L1HS, and SVA-D subfamilies.

Representing the first of its kind, our large-scale comparative genomics study has shown that mobile elements evolved quite differently among different groups and species of primates, indicating that differential DNA transposition has served as an important mechanism in primate evolution.

## INTRODUCTION

Transposable elements or mobile elements (“MEs” hereafter) are defined as genomic DNA sequences, which can change their positions or making copies and inserting into other locations in the genomes. MEs are quite abundant in genomes of higher species such as primates and plants; their contribution in the primate genomes ranges from 46.8% in the green monkey genome to 50.7% in the baboon genome (Carbone, et al. 2014; Chimpanzee Sequencing and Analysis 2005; Cordaux and Batzer 2009; Deininger, et al. 2003; Lander, et al. 2001; Locke, et al. 2011; Rhesus Macaque Genome Sequencing and Analysis, et al. 2007; Scally, et al. 2012; Yan, et al. 2011). This percentage is expected to increase slightly in these genomes due to further improvements of the genome sequences and repeat annotation, especially for the non-human primate genomes.

By the mechanism of DNA transposition, MEs can be divided into two major classes: DNA transposons and retrotransposons (Stewart, et al. 2011). DNA transposons move in the genome in a “cut and paste” style, for which they were initially called “jumping genes” (Deininger, et al. 2003). It means that they are able to excise themselves out from their original locations and move to new sites in the genome in the form of DNA, leading to no change of their copy numbers in the genome during the process (Pace Ii and Feschotte 2007). DNA transposons constituent approximately 3.6% of the primate genomes. In comparison, retrotransposons mobilize in genomes via an RNA-based duplication process called retrotransposition, in which a retrotransposon is first transcribed into RNA and then reverse transcribed into DNA as a new copy inserting into a new location in the genome (Herron 2004; Kazazian 2004). Therefore, retrotransposons move in the genome through a “copy and paste” style, which leads to an increase in their copy numbers. Retrotransposons’ high success in the primate genomes made them as the major classes of MEs, constituting on average 45% of the genomes. Depending on the presence or absence of long terminal repeats (LTRs), the retrotransposons can be further divided into LTR retrotransposons and non-LTR retrotransposons, respectively (Cordaux and Batzer 2009; Deininger, et al. 2003). In primates, the LTR retrotransposons mainly consist of endogenous retrovirus (ERVs), which are results of exogenous virus integrating into the host genomes during different stages of primate evolution (Kazazian 2004). The Short-INterspersed Elements (SINEs), the Long INterspersed Elements (LINEs), and the chimeric elements, SINE-R/VNTR/Alu (SVA), as well as processed pseudogenes, collectively represent the non-LTR retrotransposons in the primate genomes. A canonical non-LTR retrotransposon has a 3’ poly (A) tail and a pair of short repeats at the ends of the insertion sequence called target site duplications (TSDs) (Allet 1979; Grindley 1978). TSDs are a result and hallmark of the L1 driven target-<u>p</u>rimed reverse transcription (TPRT) mechanism (Anwar, et al. 2017; Goodier 2016).

MEs have been considered as “junk DNA” for nearly half of a century after McClintock first reported in the maize genome (Doolittle and Sapienza 1980; McCLINTOCK 1951). However, during the last two decades, researchers have obtained ample evidence that MEs made significant contributions to genome evolution, and they can impact gene function via a variety of mechanisms. These mechanisms include but are not limited to generation of insertional mutations and causing genomic instability, creation of new genes and splicing isoforms, exon shuffling, and alteration of gene expression and epigenetic regulation (Callinan, et al. 2005; Han, et al. 2004; Han, et al. 2007; Han, et al. 2005; Konkel and Batzer 2010; Mita and Boeke 2016; Quinn and Bubb 2014; Sen, et al. 2006; Symer, et al. 2002; Szak, et al. 2003; Wheelan, et al. 2005). MEs also contribute to genetic diseases in human via both germline and somatic insertions (Anwar, et al. 2017; Goodier 2016).

Furthermore, MEs have intimate associations with other repetitive elements such as microsatellite repeats/tandem repeats in plants (Ramsay, et al. 1999) or may have involved in the genesis of these repetitive elements (Wilder and Hollocher 2001). It was shown more recently that MEs contribute to at least 23% of all minisatellites/satellites in the human genome (Ahmed and Liang 2012). MEs have been accumulating along primate evolution. Although the majority of MEs are “fixed” in the primate genomes meaning they are shared by all primate genomes, certain MEs are uniquely owned by a particular species or lineage. A recent study has suggested that regulatory regions derived from primate and human lineage-specific MEs can be transcriptionally activated in a heterologous regulatory environment to alter histone modifications and DNA methylation, as well as expression of nearby genes in both germline and somatic cells (Ward, et al. 2013). This observation suggests that lineage- and species-specific MEs can provide novel regulatory sites in the genome, which can potentially regulate nearby genes’ expression, and ultimately lead to in lineage- and species-specific phenotypic differences. For example, it was recently shown that lineage-specific ERV elements in the primate genomes can act as IFN-inducible enhancers in mammalian immune defenses (Chuong, et al. 2016).

Past and ongoing studies on MEs in the primate genomes have been mainly focused on the human genome, examining mostly the youngest and active members that contribute to genetic variations among individuals (Battilana, et al. 2006; Ewing and Kazazian 2011; Jha, et al. 2009; Ray, et al. 2005; Seleme, et al. 2006; Stewart, et al. 2011; Wang, et al. 2006). For example, studies have shown that certain members from *L1, Alu, SVA*, and *HERV* families are still active in the human genome, and they are responsible for generating population-specific MEs (Ahmed, et al. 2013; Beck, et al. 2010; Benit, et al. 2003; Mills, et al. 2007; Wang, et al. 2005a). Besides these, limited analyses of species-specific mobile elements have also been performed in a few primate genomes. The first of such study was done by Mills and colleagues, who analyzed species-specific MEs in both the human and chimpanzee genomes based on earlier versions of the genomic sequences (GRHc35/hg17 and CGSC1.1/panTrol1.1), which led to the identification of a total of 7,786 and 2,933 MEs that are uniquely owned by human and chimpanzee, respectively (Mills, et al. 2006). However, these early studies of species-specific MEs were limited by the low quality of available genome sequences and unavailability of other primate genome sequences. Recently, we have provided a comprehensive compilation of MEs that are uniquely present in the human genomes. By making use of the most recent genome sequences for human and many other closely related primates and a robust multi-way comparative genomic approach, we identified a total of 14,870 human-specific MEs, which contribute to 14.2 Mbp net genome sequence increase (Tang, et al. 2018). Other studies focused on species-specific MEs target on either one particular ME type and/or a few primate genomes. For example, Steely and colleague have recently ascertained 28,114 baboon-specific Alu elements by comparing the genomic sequences of baboon to both rhesus macaque and human genomes (Steely, et al. 2018).

Despite these many small-scale studies, a large-scale systematic comparative analysis of DNA transposition among primates is still lacking. In this study, we adopted our robust multi-way comparative genomic approach used for identifying human-specific MEs to analyze species-specific MEs in eight primate genomes, representing the *Hominidae* family and the *Cercopithecidae* family of the primates. Our analysis identified a total of 230,855 species-specific MEs (SS-MEs) in these genomes, which collectively contribute to ∼82 Mbp genome sequences, revealing significant differential DNA transposition among primate species.

## Materials and Methods

### Sources of primate genome sequences

For our study, we chose to include four members from each of the *Hominidae* and *Cercopithecidae* primate families. All genome sequences in fasta format and the corresponding RepeatMasker annotation files were downloaded from the UCSC genomic website (http://genome.ucsc.edu) onto our local servers for in-house analysis. In all cases except for gorilla, the most recent genome versions available on the UCSC genome browser site at the time of the study were used. The four *Hominidae* genomes include the human genome (GRCh38/UCSC hg38), chimpanzee genome (May 2016, CSAC Pan_troglodytes-3.0/panTro5), gorilla genome (Dec 2014, NCBI project 31265/gorGor4.1), and orangutan genome (Jul. 2007, WUSTL version Pongo_albelii-2.0.2/ponAbe2). For gorilla genome, there is a newer version (Mar. 2016, GSMRT3/gorGor5) available, but not assigned into chromosomes, making it difficult to be used for our purpose. The four *Cercopithecidae* genomes include green monkey genome (Mar. 2014 VGC Chlorocebus_sabeus-1.1/chlSab2), crab-eating macaque genome (Jun. 2013 WashU Macaca_fascicularis_5.0/macFas5), rhesus monkey genome (Nov. 2015 BCM Mmul_8.0.1/rheMac8), and baboon (Anubis) genome (Mar. 2012 Baylor Panu_2.0/papAnu2).

### Identification of species-specific mobile element sequences (SS-MEs)

We used a computational comparative genomic approach as previously described (Tang, et al. 2018) to identify SS-MEs. In this approach, the presence/absence status of a mobile element in the orthologous regions of other genomes is determined by focusing on both whole genome alignment using liftOver and local sequence alignment using BLAT (Hinrichs, et al. 2006; Kent 2002).

#### LiftOver overchain file generation

A total of 56 liftOver chain files were needed for each of the 8 genomes used in this study. These files contain information linking the orthologous positions in a pair of genomes based on lastZ alignment (Harris 2007). Twenty-two of these were available and downloaded from the UCSC genome browser site, while the remaining 34 liftOver chain files, mostly for linking between non-human primate genomes, were generated on a local server using a modified version of UCSC pipeline RunLastzChain (http://genome.ucsc.edu).

#### Pre-processing of MEs

Our starting lists of MEs in each primate genome were those annotated using RepeatMasker. Since RepeatMasker reports fragments of MEs interrupted by other sequences and internal inversions/deletions as individual ME entries, we performed a pre-process to integrate these fragments back to ME sequences representing the original transposition events as previously described (Tang, et al. 2018). This step is critical for obtaining more accurate counting of the transposition events, and more importantly for obtaining correct flanking sequences to identify SS-MEs and their TSDs.

#### Identification of SS-MEs

As previously described (Tang, et al. 2018), our strategy for identifying SS-MEs is to examine ME insertions and the two flanking regions for each of the MEs (after integration) in a genome and compare with the sequences of the corresponding orthologous regions in all other genomes. If an ME is determined with high confidence to be absent from the orthologous regions of all examined out-group primate genomes, and then it is considered to be species-specific in this genome. Briefly, we used two tools, BLAT, and liftOver (http://genomes.ucsc.edu), for determining the orthologous sequences and the species-specific status of MEs using the aforementioned integrated RepeatMasker ME list as input. Only the MEs that are supported to be unique to a species by both tools were included in the final list of SS-MEs.

### Identification of TSDs, transductions, and insertion mediated-deletions (IMD)

The TSDs, as well as transductions and IMDs for all SS-MEs, were identified using in-house Perl scripts as described previously (Tang, et al. 2018). For those with TSDs successfully identified, a 30-bp sequence centered at each insertion site in the predicted pre-integration alleles were extracted after removing the ME sequence and one copy of the TSDs from the ME alleles. Entries with identified TSDs and extra sequences between the ME and either copy of the TSDs are considered potential candidates for ME insertion-mediated transductions and were subject to further validation as previously described. For entries without TSDs, if there are extra sequences at the pre-integration site in the out-group genomes, they were considered candidates for IMDs, which were subject to further validation.

### Identification of most recent SS-MEs

The list of SS-MEs in each genome was used to identify a subset of MEs that represent the most recent ME copies based on sequence divergence level by running all-against-all sequence alignment among all SS-MEs from a genome using BLAT. The list of human-specific MEs reported in Tang, et al. 2018 was used as a reference dataset to determine a set of optimal BLAT parameters (minScore >=100; minIdentity >= 98) that identifies 95% of the SS-MEs in the human genome as the most recent MEs. These criteria were then applied to identifying the most recent SS-MEs in each of the other genomes.

### Analysis of SS-MEs’ association with genes in the primate genomes

The non-human primate genomes used in this study are not as well annotated as the human genome. Therefore, we used the genomic coordinates of genes in the human genome breaking down to individual exons based on GENCODE gene annotation (Harrow, et al. 2012) and NCBI RefSeq data (Pruitt, et al. 2007). The entire human genome sequences were divided into a non-redundant list of categorized regions in gene context, including coding sequence (CDS), non-coding RNA, 5’-UTR, 3’-UTR, promoter (1kb), intron, and intergenic regions using an in-house Perl script as previously described (Tang, et al. 2018). This order of genic region categories as listed above was used to set the priority from high to low in handling overlapping regions between splice forms of the same gene or different genes. For example, if a region is a CDS for one transcript/gene and is a UTR or intron for another, then this region would be categorized as CDS. The corresponding orthologous regions of these regions in each of the non-human primate genomes were identified using the liftOver tool.

### Statistical analysis

The statistical analysis and figure plotting were performed using a combination of Linux shell scripts, R and Microsoft Excel.

## RESULTS

### he differential ME profiles of primate genomes

We first compared the profiles of all repeat elements across the eight primate genomes. As shown in fig. S1 and Tables S1 & S2, the total percentage of all repeat elements in the genomes ranges from ∼49% (green monkey) to ∼57% (baboon) (fig. S1) with MEs representing the majority of the repeats, contributing to ∼46.8% (green monkey) to 50.9% of the genomes (baboon), averaging at 48.5% (Table S1). Among the non-ME repeats (low complexity, rolling circle, RNA, satellite, and simple repeat), which collectively contribute 2% to 6% of the genomes, the satellites show the largest degree of variations among the eight genomes ranging from 0.2% in rhesus to 4.3% in the baboon genome (Table S2). For both the total repeat content and ME content, the baboon genome has the highest percentages (56.8% for all repeats and 50.7% for MEs) and the green monkey genome has the lowest percentages (48.8% for all repeat and 46.8% for MEs).

The percentages of MEs by the major ME class in a genome is quite similar across genomes with a few exceptions (fig. S2A). DNA transposons account for 3.5% to 3.8% (3.6% on average) in the genomes, while retrotransposons are much more successful, contributing to 43.3% to 47.1% (48.5% on average) of the genomes and serving as the main contributors for the overall ME differences among genomes (fig. S2A & Table S1). Within the retrotransposon group, LINE, SINE, and LTR are the most successful classes, contributing to an average 21.8%, 13.9% and 9.1% of the primate genomes by sequence size on average, respectively (Table S1). The profiles of SVAs are quite different between the *Hominida*e family and *Cercopithecidae* family with the copy number ranging from a few hundreds (∼0.002%) in the latter group to a few thousand copies (0.1%) the former group (fig. S2B & Table 1). We also compared other attributes of MEs, such as copy number and average length. As shown in Table 1, the SINEs are the most successful MEs by copy number (1,652,055 on average), ranging from 1,602,634 (orangutan) to 1,706,611 (rhesus). On average, there are 930,532 LINEs, 477,258 LTRs, and 382,247 DNA transposons per genome (Table 1). The DNA transposons and SINEs have the shortest average length at 259 bp and 231 bp, respectively, and the LINEs and LTRs average at 644 bp and 526 bp, respectively (Table S3). While similar ME classes share a similar average length across genomes in most cases, SVAs in the *Hominidae* family are quite different from the MacSVAs in the *Cercopithecidae* family, with the average lengths being ∼900 bp for SVAs and ∼350 bp for MacSVAs (Table S3). Furthermore, within the *Hominidae* family, the average lengths of SVAs in the orangutan and gorilla genomes (1,162 bp and 547 bp, respectively) are quite different than that of SVAs in human and chimpanzee genomes (856bp and 870bp, respectively). The subfamilies of SVAs with the longest average length are different across genomes with it being SVA-D for both human and chimpanzee, SVA-A and SVA-B for orangutan, and SVA-C for gorilla, which is significantly shorter in length than SVA-C and the longest SVA subfamilies in the other *Hominida*genome genomes (Table S4). Notably, the SVA-E and SVA-F subfamilies are only found in the human genome (Table S4).

**Table 1.** Mobile element (ME) compositions by copy number in eight primate genomes.

| Genome | Human |  | Chimpanzee |  | Gorilla |  | Orangutan |  | Green monkey |  | Crab-eating macaque |  | Rhesus |  | Baboon |  |
| --- | --- | --- | --- | --- | --- | --- | --- | --- | --- | --- | --- | --- | --- | --- | --- | --- |
| ME class | raw counts | integrated counts | raw counts | integrated counts | raw counts | integrated counts | raw counts | integrated counts | raw counts | integrated counts | raw counts | integrated counts | raw counts | integrated counts | raw counts | integrated counts |
| SINE | 1,779,233 | 1,689,416 | 1,771,039 | 1,682,623 | 1,722,434 | 1,638,587 | 1,689,629 | 1,602,634 | 1,709,337 | 1,616,578 | 1,721,680 | 1,631,626 | 1,796,021 | 1,706,611 | 1,801,595 | 1,648,361 |
| LINE | 1,516,226 | 969,873 | 1,551,601 | 1,000,667 | 1,533,883 | 1,000,110 | 1,428,157 | 907,077 | 1,426,343 | 886,492 | 1,414,592 | 875,720 | 1,477,648 | 948,851 | 1,471,152 | 899,503 |
| SVA | 5,397 | 4,933 | 5,358 | 4,931 | 5,492 | 4,809 | 2,771 | 2,328 | 145 | 133 | 145 | 130 | 152 | 137 | 169 | 141 |
| LTR | 720,177 | 496,946 | 723,412 | 499,454 | 707,051 | 494,156 | 671,620 | 470,734 | 676,130 | 462,936 | 664,942 | 460,094 | 695,510 | 480,535 | 701,611 | 467,533 |
| DNA | 483,994 | 399,590 | 510,250 | 421,580 | 503,480 | 418,454 | 429,467 | 347,471 | 445,724 | 361,048 | 443,909 | 359,802 | 486,991 | 401,546 | 459,662 | 369,684 |
| **Total** | **4,505,027** | **3,560,758** | **4,561,660** | **3,609,255** | **4,472,340** | **3,556,116** | **4,221,644** | **3,330,244** | **4,257,679** | **3,327,187** | **4,245,268** | **3,327,372** | **4,456,322** | **3,537,680** | **4,434,189** | **3,385,222** |

### The impact of ME sequence consolidation

The initial ME lists used in this study were based on the RepeatMasker annotations obtained from the UCSC Genome Browser, and we performed integration of fragmented MEs to represent original transposition events to improve the accuracy in identifying SS-MEs and the TSDs. As shown in Table 1, the consolidation led to an average reduction of 940,000 per genome in ME counts. The eight primate genomes contain 3,454,229 MEs/genome on average after consolidation (vs. raw counts at 4,379,090/genome), with the chimpanzee genome having the largest number of MEs (3,609,255) and the green monkey genome and crab-eating macaque having the least number of MEs at 3,327,187 and 3,327,372, respectively (Table 1). The degree of integration was assessed in the degree of increase in the rate of full-length MEs (defined as ≧90% of the consensus sequence) and in average length. As shown in Table S1, the rates of full-length MEs for all classes increased after integration with LINEs showing the largest degree of increase (≧2 folds), indicating that LINEs were most frequently interrupted by post-insertion events, likely due to their longer lengths and overall older age. The relatively larger increase for LTRs than the rest ME classes other than LINEs is likely in part due to RepeatMasker’s practice of reporting internal sequences and the terminal repeats as separate entries. Full-length entries increased from 21.0% to 29.9% for DNA transposons and from 48.4% to 51.8% for SINEs in the human genome, while SVAs showed the least amount of full-length rate increase indicating their youngest age among all ME classes. Notably, SVAs in gorilla were shown to have a much lower full-length rate (11.4%) compared to the other three *Hominidae* genomes (32.8% to 36.4%) (Table S3), and this might be a result of a higher percentage of unsequenced regions in this version of the gorilla genome sequences (gorGor4). This is consistent with their average length also being the shortest among the *Hominidae* family (Table S4). The extremely low rates of full-length entries for the LINEs (0.8 to 1.4% after integration) compared to these of other ME classes, which are more than 20% (Table S3), is likely a result of heavily truncation during insertion due to their long full-length consensus sequences. As shown in Table S3, the general trend for the degree of increase from integration showed a positive correlation with the length of the ME consensus sequences, i.e., LINEs showing the largest degree of increase, while SINEs showing the least degree of increase. However, DNA transposons seem to be out of this trend by being about the same sizes as SINEs, but with a much higher degree of increases than SINEs, likely due to their relative older age. For the same reason, SVAs showed less size increase than DNA transposons despite being larger in size, likely due to their much younger ages.

### Differential level of species-specific MEs (SS-MEs) in primate genomes

To assess the level of differential DNA transposition among the primate genomes, we first examined all SS-MEs that are defined as being uniquely present in each of the examined genomes. Our analysis of SS-MEs was based on the consolidated ME lists as discussed in the previous section and it was performed using a multi-way comparative genomics approach extended from our previously described method in identifying human-specific MEs (Tang, et al. 2018). By comparing each of the either primate eight genomes to the rest seven genomes, we identified a total of 230,855 SS-MEs, consisting of 150,260 SINEs, 61,216 LINEs, 5,230 SVAs, 11,744 LTRs and 2,405 DNA transposons (Table 2).

**Table 2.** The number of species-specific mobile elements (SS-MEs) and most recent SS-MEs in eight primate genomes

| Genome | Human |  | Chimpanzee |  | Gorilla |  | Orangutan |  | Green monkey |  | Crab-eating macaque |  | Rhesus |  | Baboon |  | Total |  |
| --- | --- | --- | --- | --- | --- | --- | --- | --- | --- | --- | --- | --- | --- | --- | --- | --- | --- | --- |
| ME class | SS-ME | most recent SS-ME | SS-ME | recent SS-ME | SS-ME | recent SS-ME | SS-ME | most recent SS-ME | SS-ME | most recent SS-ME | SS-ME | most recent SS-ME | SS-ME | most recent SS-ME | SS-ME | most recent SS-ME | SS-ME | most recent SS-ME |
| SINE | 8,844 | 8,138 | 10,612 | 3,587 | 6,324 | 3,882 | 9,630 | 1,205 | 34,277 | 23,582 | 2,257 | 1,190 | 22,069 | 11,646 | 56,247 | 51,065 | 150,260 | **104,295** |
| LINE | 3,946 | 3,197 | 7,288 | 4,430 | 4,085 | 2,977 | 21,711 | 16,228 | 11,981 | 8,381 | 782 | 566 | 3,016 | 1,538 | 8,407 | 7,032 | 61,216 | **44,349** |
| SVA | 1,571 | 1,533 | 1,597 | 1,218 | 877 | 798 | 1,180 | 752 | 3 | 0 | 0 | 0 | 2 | 0 | 0 | 0 | 5,230 | **4,301** |
| LTR | 530 | 292 | 1,924 | 441 | 689 | 310 | 2,933 | 890 | 2,324 | 1,028 | 234 | 151 | 1,346 | 353 | 1,764 | 1,171 | 11,744 | **4,636** |
| DNA | 56 | 11 | 666 | 50 | 151 | 62 | 668 | 125 | 322 | 35 | 8 | 1 | 374 | 23 | 160 | 33 | 2,405 | **340** |
| **All SS-MEs** | **14,947** | **13,171** | **22,087** | **9,726** | **12,126** | **8,029** | **36,122** | **19,200** | **48,907** | **33,026** | **3,281** | **1,908** | **26,807** | **13,560** | **66,578** | **59,301** | **230,855** | **157,921** |

As shown in fig. 1 and Table 2, the copy numbers of SS-MEs are drastically different across the eight primate genomes with the baboon genome having the largest number (66,578), which is more than 20 times higher than that of the crab-eating macaque genome, which has the least number of SS-MEs (3,281). The extremely low level of SS-MEs in crat-eating macaque genome seems to be very striking by being merely one-tenth of the genome average (3,281/28,857). Furthermore, the specific compositions of SS-MEs by ME class also differ significantly across genomes with SS-SINEs represent the largest class of SS-MEs in all genomes except for orangutan genome. In the *Hominidae* genomes, the numbers of SS-SINEs are at least two times more than the number of SS-LINEs, and this difference is much larger in the *Cercopithecidae* genomes, with the numbers of SS-SINEs being 3 to 7 folds of SS-LINEs (Table 2). The orangutan genome is very unique in this aspect by having the number of SS-LINEs being more than two times of the number for SS-SINEs (Table 2). For SS-LTRs, the crab-eating macaque genome has the least number (234), followed by the human genome (530), while orangutan genome has the largest number (2,933), which is more than 10 times higher than that in the crab-eating macaque, followed by gorilla genome (2,324), then by baboon genome (1,764) and the rhesus genome (1,346). For SS-DNAs, the numbers are much smaller than all other SS-ME classes with the chimpanzee genome having the largest number (666) and the crab-eating macaque genome has the least (8), while the number in other genomes range from 56 (human) to 374 (rhesus). For SS-SVAs, the human and chimpanzee genomes have ∼1,500, while the numbers for orangutan and gorilla genomes are 1,180 and 877, respectively. While between 100 and 200 MacSVAs are present in the *Cercopithecidae* genomes, no more than 3 or zero SS-MacSVAs are detected.

**Fig. 1.**
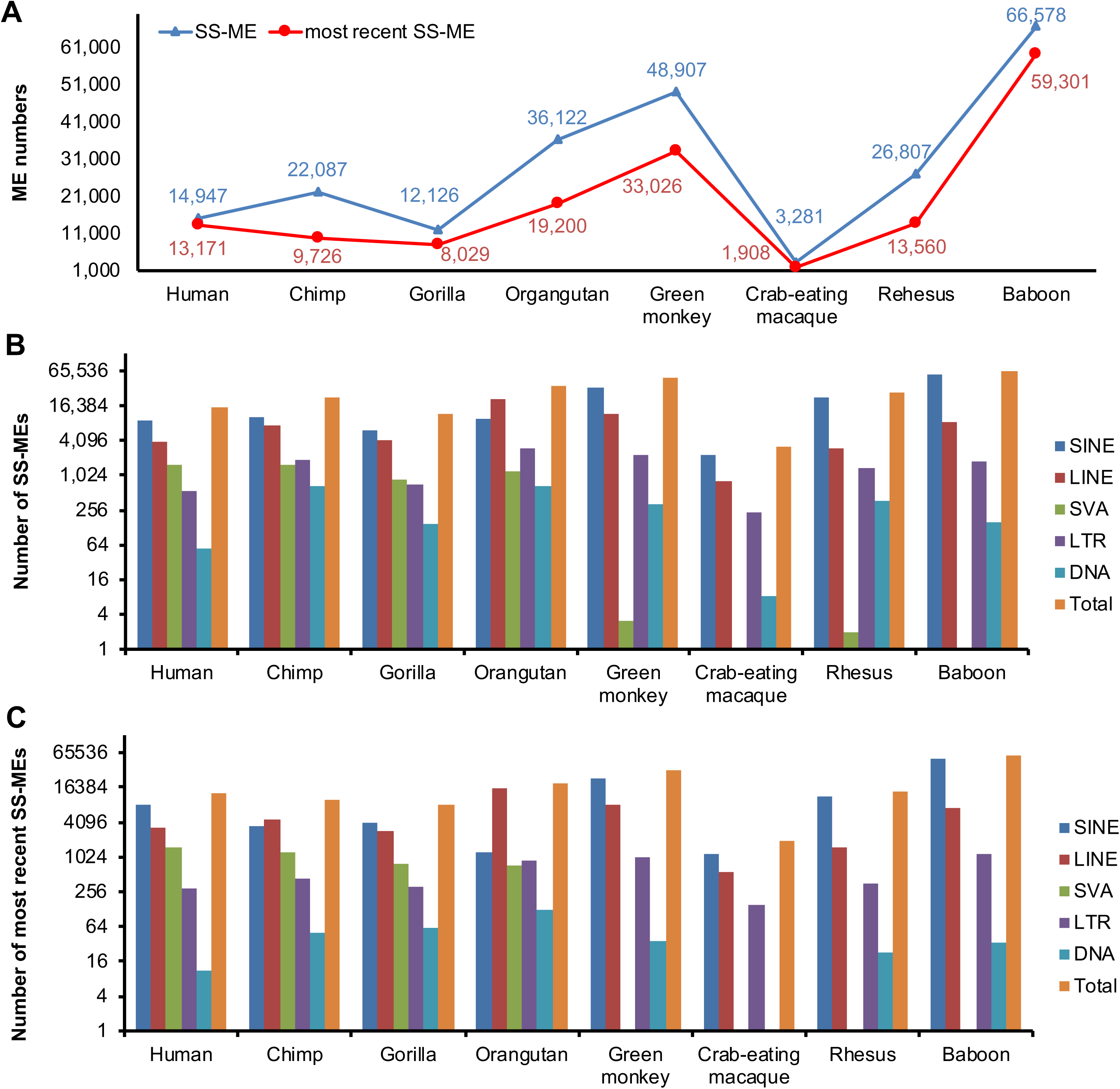
The numbers of species-specific mobile elements (SS-MEs) and most recent SS-MEs in eight primate genomes. A, The total numbers of SS-MEs and most recent SS-MEs in the primate genomes; B, The numbers of SS-MEs by ME class in the primate genomes; C, The numbers of most recent SS-MEs by ME class in the primate genomes. For both B & C, the Y-axis is in 2-based log scale.

Between the two primate families, there also seems to have a clear difference in their SS-ME profiles with the *Cercopithecidae* family having ∼3.2 times SS-SINEs than the *Hominidae* family (28,713/genome vs. 8,853/genome), but a lower number of SS-LINEs (4,379/genome) than the *Hominidae* family (9,258/genome), leading to an overall higher of SS-MEs than the latter (36,393/genome vs. 21,321/genome) (Table S5). While the level of DNA transposition seems to be more or less similar (within 1 order of differences) among the *Hominidae* genomes as measured by the total number of SS-MEs, it differs dramatically (more than 1 order) among the members of the *Cercopithecidae* family by having members both with the lowest and highest number of SS-MEs among all eight genomes in the same family.

In addition to comparison by copy numbers, we have also examined the SS-ME profiles by normalizing SS-MEs as the percentage of all SS-MEs for each class in the genomes (fig. 2A). On average, SS-LINEs contributed to ∼53.5%, and SS-SINEs contributed to ∼31.7% of all SS-MEs in the genomes, while SS-LTRs, SS-SVAs, and SS-DNAs account for ∼9.1%, ∼5.3%, and ∼0.4%, respectively. Similar to SS-ME composition by copy number, the composition of SS-MEs by ME class in sequence length are also quite different between the *Hominidae* family and *Cercopithecidae* family. For the *Hominidae* family, the top contributors are LINEs, contributing to an average of ∼68.0% of the total SS-MEs in a genome, while for the *Cercopithecidae* family, the top contributors are SINEs (average at 53.9%) in three of the four genomes due to their high numbers, despite their shorter lengths than LINEs (fig. 2B).

**Fig. 2.**
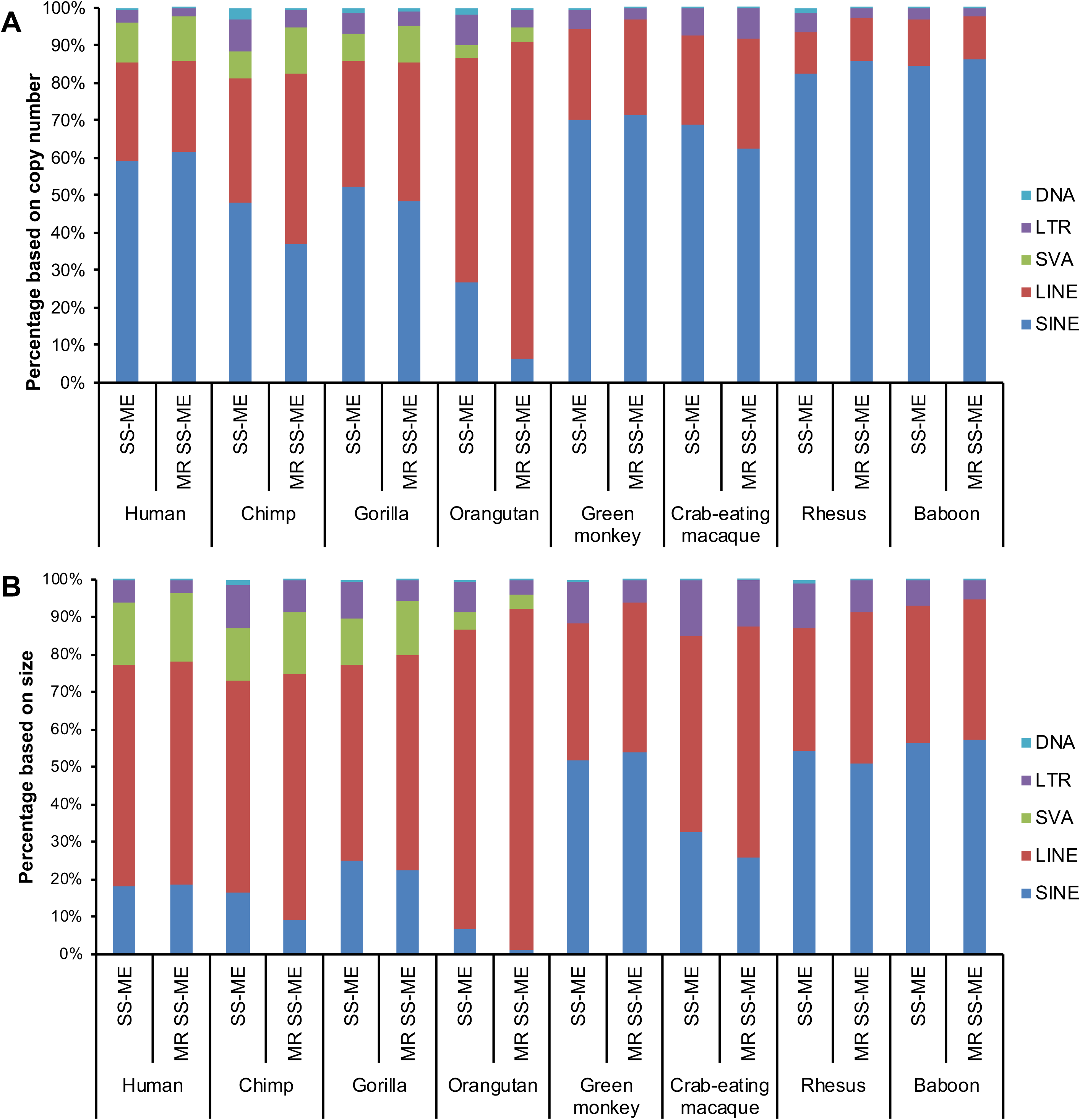
The compositions of species-specific mobile elements (SS-MEs) and most recent SS-MEs (MR SS-MEs) by ME class in eight primate genomes. A, The percentage of each SS-ME and most recent SS-ME by ME class based on copy numbers; B, The percentage of each SS-ME and most recent SS-ME by ME class based on total sequence length.

### Differential level of most recent SS-MEs in primate genomes

Since the number of SS-MEs identified in each genome is directly impacted by its evolutionary distance to the next closest genome among the genomes examined, it might not be the best comparable measure of the recent DNA transposition activity level in these genomes. For this reason, we also collected a subset of SS-MEs, which were involved in most recent transposition events seen as sharing a very high level of sequence similarity (≥98% identify over 100 bp) with another SS-ME copy in the same genome. Since the same criteria were applied to all genomes, the numbers of these most recent SS-MEs can be used to measure and compare recent DNA transposition activity across genomes.

As shown in Table 2 and fig. 1A, the overall trend for the number of most recent SS-MEs among the genomes is similar to that of SS-MEs with the baboon genome having the highest number of most recent SS-MEs (59,301) and the crab-eating macaque genome having the lowest number (1,908) (fig. 1A) and the composition by ME class also being mostly similar between the two sets of SS-MEs for all genomes (fig. 1B & C). When compared between the primate families, the *Hominidae* has a lower average number of most recent SS-MEs per genome than *Cercopithecidae* (∼12,500 vs. ∼27,000), similar to the SS-ME profile (Table S5). The fact that the crab-eating macaque genome has the lowest number of most recent SS-MEs, as in the case of SS-MEs reinforces an extremely low level of DNA transposition activity in this genome (fig. 1A). Interestingly, the orangutan genome has 16,228 most recent SS-LINEs, being the highest among all eight genomes or ∼2 times higher the 2^nd^ highest, the green monkey genome (8,381) or ∼40 times higher than that of the crab-eating macaque genome (566) (Table 2). On the other hand, the orangutan genome has a very low number of most recent SS-SINEs (1,205), very close the 1,190 SS-SINEs in the crab-eating macaque genome (Table 2). By the ratio of SINEs/LINEs in the same genome, the orangutan genome is very low for SS-MEs (0.44) and even much lower for the most recent SS-MEs (0.07) compared to other genomes, which have this ratio being ∼7 for rhesus and baboon genomes and ∼2.5 for human, green, and crab-eating macaque genomes for both SS-MEs and most recent SS-MEs. The chimpanzee genome has lower than average ratio for SS-MEs (∼1.5) and much lower but not the lowest (0.8) for most recent SS-MEs (direct ratio values based on data in Table 2 not shown).

Among the eight genomes, the ratio of most recent SS-MEs within the SS-MEs ranges from 0.44 to 0.89 with the chimpanzee genome being the lowest and the baboon genome being the highest (fig. 3A). Among the remaining genomes, the human genome (0.88) is very close to the baboon genome (0.89), while the rest 5 genomes have a ratio between 0.51 to 0.68 (fig. 3A). This profile of the ratio of the most recent SS-MEs by ME class are quite different across the eight genomes with the human genome showing the highest ratio for all three non-LTR retrotransposon classes (LINE, SINE, and SVA), the baboon genome also showing a low ratio for SINE, LINE, and LTR, and the crab-eating genome having the lowest or 2^nd^ lowest ratios for most ME classes (fig. 3B & C). The profile represented by the ratios of most recent SS-MEs by ME class are also very different across the genomes (fig. 3C). LINEs have more consistent high ratios of most recent SS-MEs among genomes, while SINEs have a high level of variability across the genomes ranging from 0.13 (orangutan) to 0.92 (human) (fig. 3C). DNA transposons show a low ratio of most recent SS-MEs in most genomes, being the lowest overall among the ME classes, but with the gorilla genome standing out by having a ratio that is several times higher than average (0.41 vs. 0.17) (fig. 3C).

**Fig. 3.**
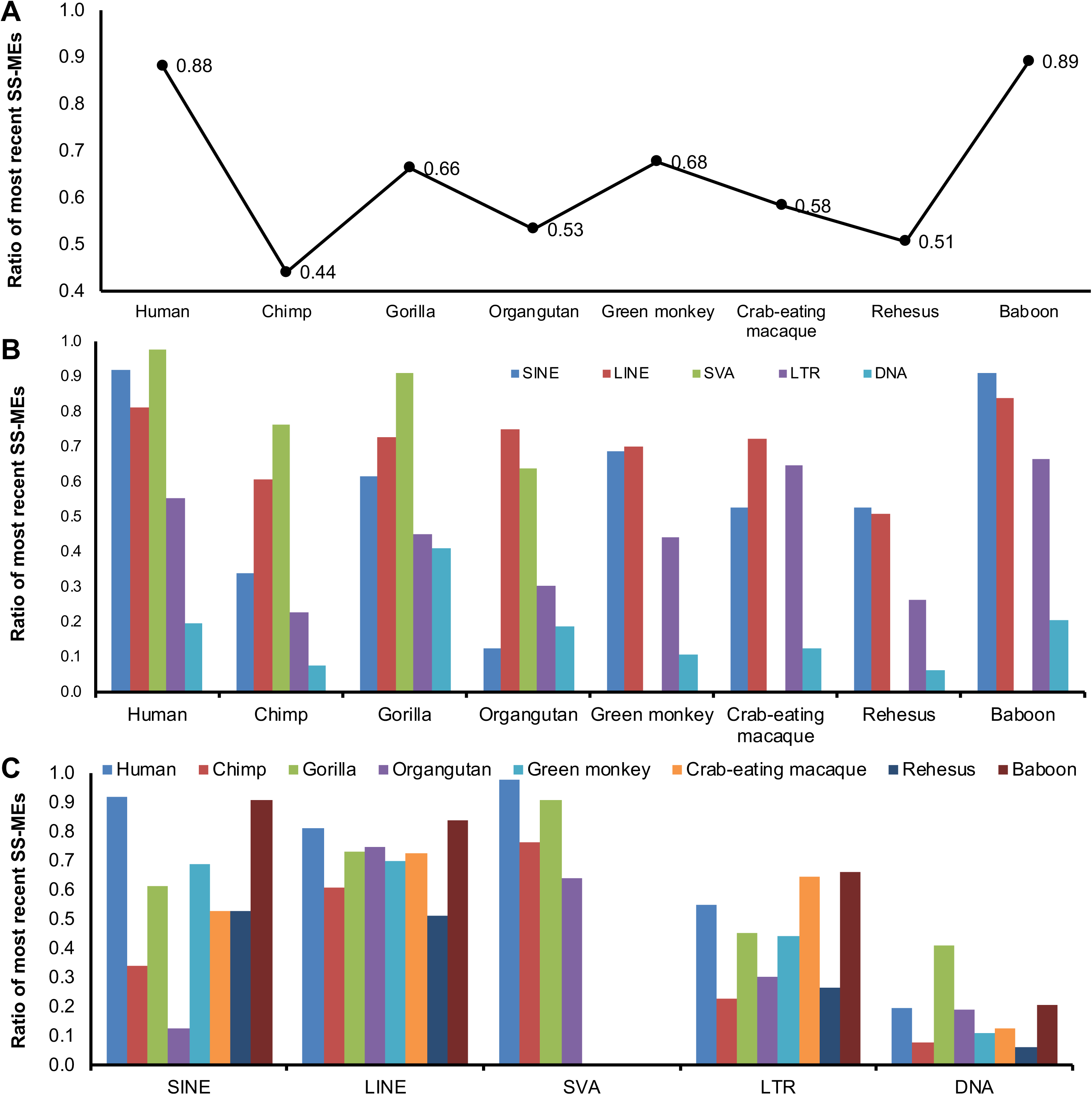
The ratios of most recent species-specific mobile elements (SS-MEs) in the primate genomes. The ratio is calculated by dividing the number of the most recent SS-MEs in a genome by the number of SS-MEs in a genome. A, The ratios for the total number of most recent SS-MEs in the genomes; B, The ratios of most recent SS-MEs broken down into ME classes grouped for each genome; C, The ratios of most recent SS-MEs broken down to genomes for each ME class.

It is very interesting to notice that between the human and chimpanzee genomes, representing probably the two most closely related genomes among the eight primate genomes analyzed, while the chimpanzee genome has a much larger number of SS-MEs (22,087 vs. 14,947 in the human genome), the human genome has a much larger number of most recent SS-MEs (13,171 vs 9,726 in the chimpanzee genome) (Table 2 and fig. 1A). To better understand these significant differences, we analyzed the age and activity profiles of SS-MEs by ME class in these two genomes based on sequence divergence level of SS-MEs. As shown in Fig. 4, the age profiles of individual SS-ME classes are quite different between the two genomes. The human genome showed a lower level of overall activity early on (fig. 4A) but a much more rapid increase of activity recently as reflected by the number of SS-MEs at high sequence similarity (fig. 4A to 4G). The higher most recent DNA transposition activity in the human genome is seen for SINEs and SVAs (fig. 4C & E) with SINEs contributing most to the higher number of most recent SS-MEs in the genome compared to the chimpanzee genome. The chimpanzee genome showed a higher most recent activity for LINEs, LTRs, and DNA transposons (fig. 4D/F/G), but the degree of differences is much smaller than that for SINEs, leading to a lower number of most recent SS-MEs than in the human genome. Interestingly, SVAs in the human genome showed a lower activity early on, but a quicker acceleration, followed by a trend of plateau or even a slightly lower towards the most recent period (fig. 4E). In contrast, SVAs in chimpanzee genome showed mostly lower (than in the human genome) but steady increase of activity all the way to the most recent period (fig. 4E). This seems to correlate well with the higher level of SVA-D activity in the human genome than in the chimpanzee genome (fig. 5).

**Fig. 4.**
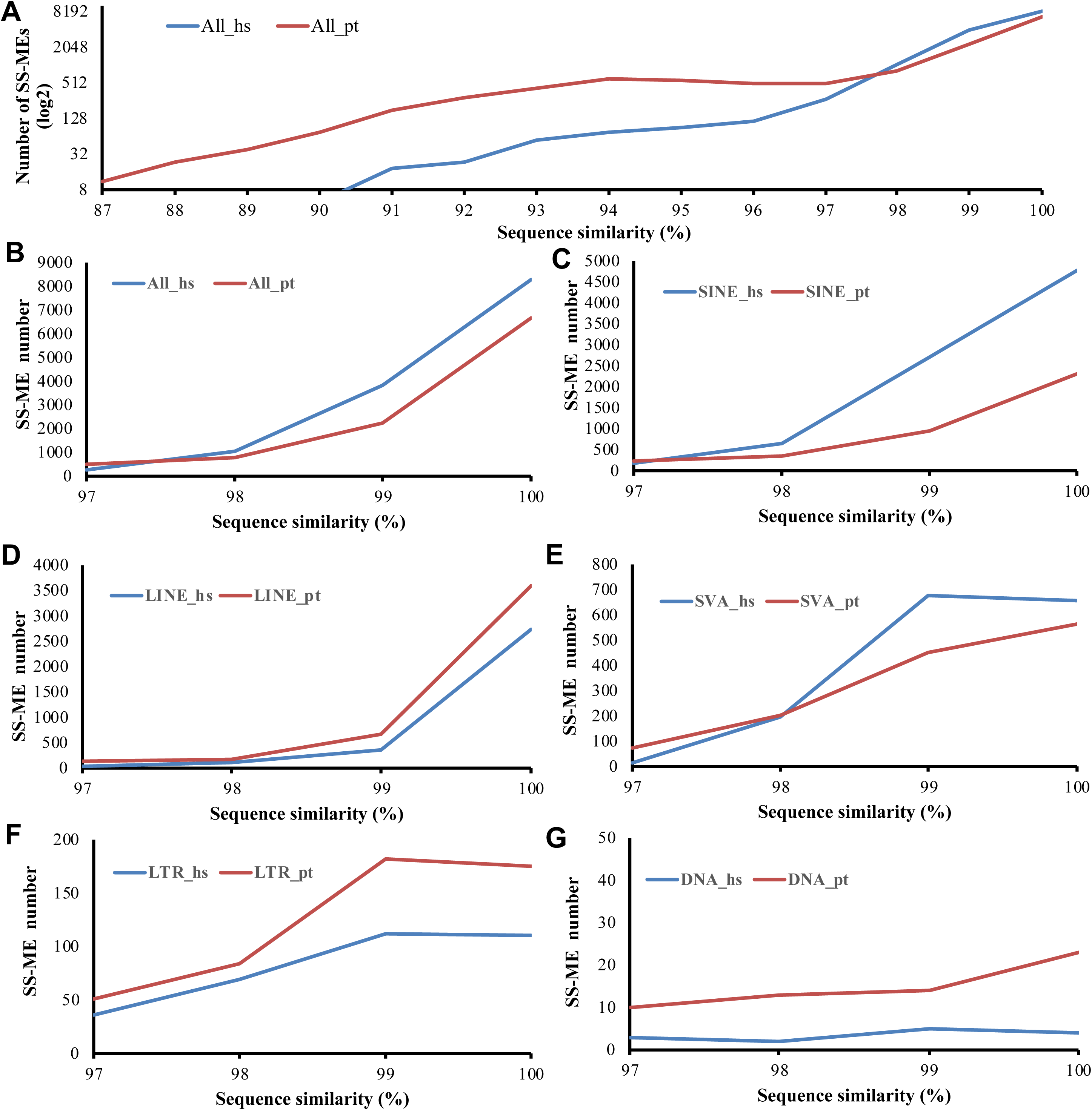
The activity profiles of more recent specifies-specific mobile elements (SS-MEs) by ME class in the human and chimpanzee genomes. The numbers of SS-MEs with sequence similarity at 87% (A) or and 97% (B-E) or more to another copy of SS-MEs in the human (“_hs”) and chimpanzee (“_pt”) genomes are shown for each ME class. A & B. all SS-MEs combined; C. SS-SINEs; D. SS-LINE; E. SS-SVAs; F. SS-LTRs; G. SS-DNAs.

**Fig. 5.**
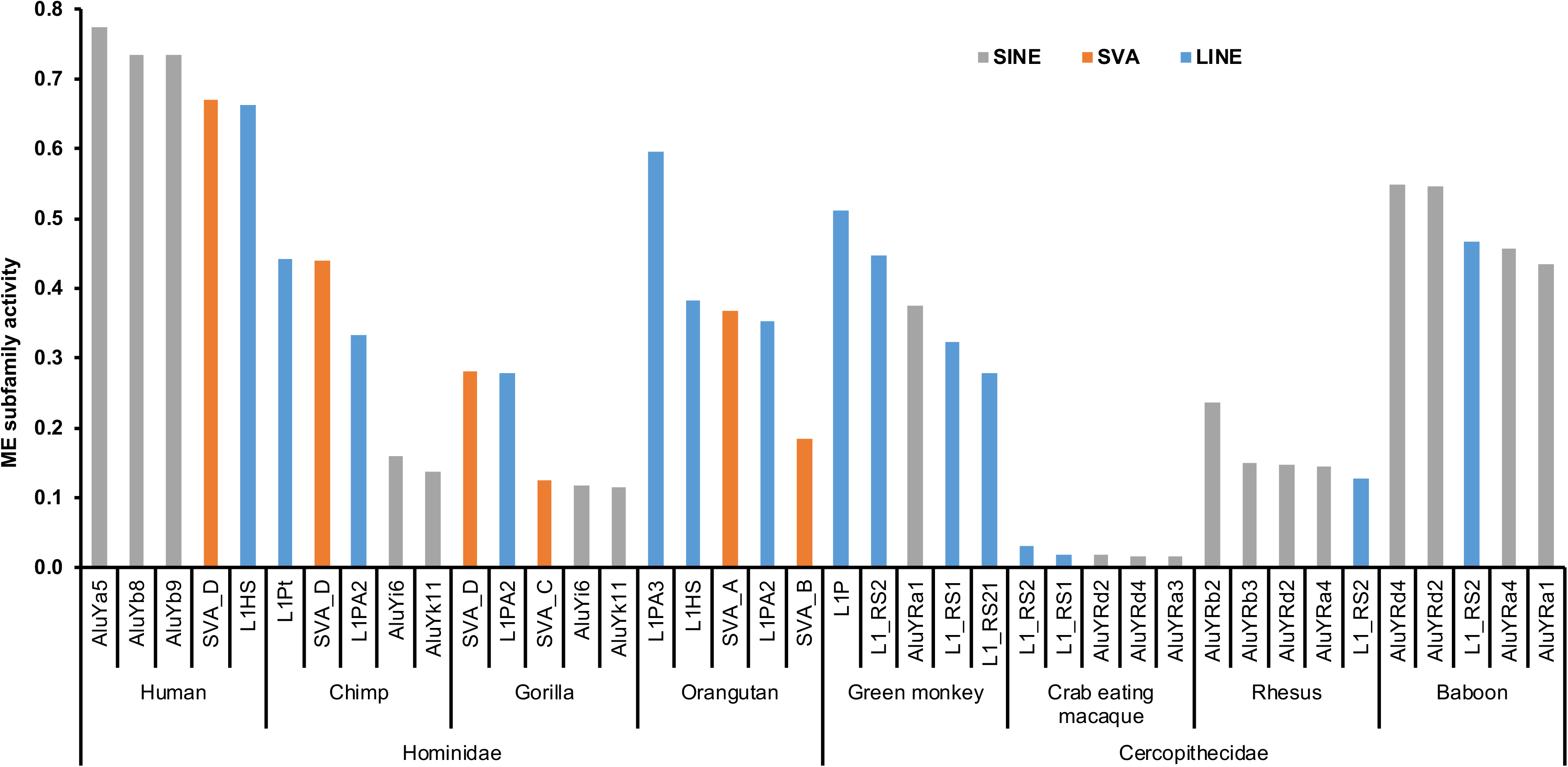
Most active subfamilies of mobile elements (MEs) in the eight primate genomes. The top 5 active ME subfamilies in each primate genome are listed. The activity level of each ME subfamily was calculated by dividing the numbers of most recent SS-MEs with the total numbers of MEs in the subfamily.

### The most active ME subfamilies in the eight primate genomes

The lists of most recent SS-MEs provides an unbiased measure for the relative level of DNA transposition activity across the genomes, as well as among different ME classes and subfamilies. Table 3 shows the most recent transposition activity by ME class in each genome calculated as the percentage of the most recent SS-MEs in all MEs in the class. At the ME class level, for SINEs, the baboon genome has the highest activity (3.1%), which is more than double of the 2^nd^ highest (green monkey, 1.46%) and more than 40 times higher than the lowest (crab-eating macaque, 0.07%) (Table 3). For LINEs, the orangutan genome has the highest activity (1.79%), which is approximately two times of that for the 2^nd^ highest (green monkey, 0.95%) and 30 times higher than the lowest (crab-eating macaque, 0.06%). For SVAs, LTRs, and DNA transposons, the most active genomes are human/orangutan, baboon, and orangutan, respectively (Table 3). As expected, SVAs, being the youngest ME class, have the highest activity among all ME classes. For example, in the human genome, the activity of SVAs (31%) is ∼65 times higher than that of SINEs (0.48%), being the second most active ME class in the genome (Table 3).

**Table 3.** Recent DNA transition activity in the eight primate genomes.

| Table 3. Recent DNA transition activity in the eight primate genomes. |  |  |  |  |  |  |  |  |
| --- | --- | --- | --- | --- | --- | --- | --- | --- |
| ME class | Human | Chimpanzee | Gorilla | Orangutan | Green monkey | Crab-eating macaque | Rhesus | Baboon |
| SINE | 0.48* | 0.21 | 0.24 | 0.08 | 1.46 | 0.07 | 0.68 | 3.10 |
| LINE | 0.33 | 0.44 | 0.30 | 1.79 | 0.95 | 0.06 | 0.16 | 0.78 |
| SVA | 31.08 | 24.70 | 16.59 | 32.30 | 0 | 0 | 0 | 0 |
| LTR | 0.06 | 0.09 | 0.06 | 0.19 | 0.22 | 0.03 | 0.07 | 0.25 |
| DNA | 0.00 | 0.01 | 0.01 | 0.04 | 0.01 | 0.00 | 0.01 | 0.01 |

\*, the percentage of species-specific mobile elements (SS-MEs) in a ME class in the genome

Further details in the activity profiles were revealed by examining the most active ME subfamilies in each genome. Fig. 5 shows the top 5 ME subfamilies and their relative activity level in each genome, indicating that each genome has a unique profile of active MEs that differ not only by ME subfamilies but also by their relative levels of activity. In the human genome, AluYa5, AluYb8/9 are the most active SINE subfamilies; L1HS, L1PA3, and L1P are the most active LINE subfamilies; SVA_D is the most active SVA subfamily. The activity levels of these 5 subfamilies are higher than any other ME subfamilies in any other genomes, indicating that the human genome has the highest most recent DNA transposition activity among the eight genomes. In contrast, the crab-eating macaque genome lacks a single highly active ME subfamily (Table S6, fig. 5 & S3). Interestingly, the baboon genome, which has the highest number of SS-MEs and most recent SS-MEs, as well as the highest ratio of most recent SS-MEs (fig. 1A & 2A), has no single ME subfamily being extremely active as seen in the human genome, but all top 5 ME subfamilies are from SINE and with similarly high activity levels (fig. 5), making the baboon genome as the most active genome for SINE transposition. Three of the top 5 ME subfamilies in the orangutan genomes are from LINEs, making it the most active genome for LINE transposition. Between the two groups of the primates, the *Hominidae* family has AluYa/b subfamilies as the most active SINEs and *L1P* subfamilies as the most active LINEs, while the *Cercopithecidae* family has *AluYRs* as the most active SINEs and *L1_RS* subfamilies as the most active LINEs (fig. 5).

### Differential impact of DNA transposition on primate genome sizes

We compared across the eight genomes the impact SS-MEs on genome size via insertion of MEs and generation of TSDs and transductions, as well as possible genome size reduction through insertion-mediated deletions (IMD) of flanking sequences. As shown in Table 4, in all eight genomes, SS-MEs have led to a net genome size increase. Collectively, SS-MEs have contributed to a combined ∼82.3 Mbp increase in the eight genomes or on average ∼10 Mbp per genome. However, the degree of size increase varies significantly among the genomes with the orangutan genome gaining the largest increase (∼26 Mb) and the crab-eating macaque genome gaining the least (∼1.2 Mb), which is directly correlated with the overall levels of SS-MEs. Among the different types of size impact, the insertion of ME sequences is responsible for the majority of the size increase as expected, followed by IMD, transductions, and TSDs (Table 4). The orangutan genome had the largest increase from ME insertions (33.9 Mbp), which is ∼6 times higher than the lowest in the *Hominidae* family, which is the gorilla genome (∼5.8 Mbp) and ∼18 time higher than that of the crab-eating macaque genome (∼1.9 Mbp) as the lowest among all. This seems to be contributed to the highest portion of SS-LINEs in the orangutan genome. The baboon genome has the largest increase from TSDs and transductions, likely due to the largest number of SS-MEs, while the green monkey genome has the largest genome loss from IMD (Table 4). The size impact for the most recent SS-MEs is relatively more or less similar to that of SS-MEs, except that the baboon genome replaces the orangutan genome as having the largest size increase from ME insertions due to its high ratio of most recent SS-MEs. For the genomes of chimpanzee, gorilla, green monkey, and rhesus, the most recent SS-MEs led to more genome size increase than all SS-MEs due to lower portions of insertion-mediated deletions (IMDs) among most recent SS-MEs (Table 4).

**Table 4.** Impact of species-specific mobile elements (SS-MEs) and most recent SS-MEs on genome size (Kb)

|  |  |  |  |  |  |  |  |  |  |  |  |  |  |  |  |  |  |  |  |  |
| --- | --- | --- | --- | --- | --- | --- | --- | --- | --- | --- | --- | --- | --- | --- | --- | --- | --- | --- | --- | --- |
| Genome | Human |  | Chimpanzee |  | Gorilla |  | Orangutan |  | Green monkey |  | Crab-eating macaque |  | Rhesus |  | Baboon |  | Total |  | Average |  |
| Sequence change type* | SS-ME | most recent SS-ME | SS-ME | most recent SS-ME | SS-ME | most recent SS-ME | SS-ME | most recent SS-ME | SS-ME | most recent SS-ME | SS-ME | most recent SS-ME | SS-ME | most recent SS-ME | SS-ME | most recent SS-ME | SS-ME | most recent SS-ME | SS-ME | most recent SS-ME |
| ME insertion | 14,259 | 13,170 | 16,274 | 11,073 | 5,895 | 4,842 | 33,924 | 26,696 | 17,330 | 12,390 | 1,797 | 1,389 | 11,074 | 6,741 | 29,342 | 26,784 | 129,894 | 103,085 | 16,237 | 10,900 |
| TSD | 171 | 161 | 118 | 94 | 89 | 73 | 243 | 164 | 353 | 274 | 15 | 11 | 139 | 107 | 581 | 543 | 1,709 | 1,427 | 214 | 126 |
| Transduction | 687 | 552 | 1,033 | 664 | 1,086 | 668 | 3,741 | 2,403 | 4,435 | 2,870 | 646 | 371 | 2,616 | 1,731 | 6,063 | 5,381 | 20,307 | 14,640 | 2,538 | 1,323 |
| IMD | -977 | -494 | -11,403 | -1,878 | -4,073 | -1,928 | -12,381 | -4,281 | ##### | -7,276 | ##### | -603 | -10,700 | -3,156 | -12,448 | -9,476 | -69,543 | -29,092 | -8,693 | -2,802 |
| Net total | 14,141 | 13,390 | 6,021 | 9,953 | 2,996 | 3,655 | 25,527 | 24,982 | 5,742 | 8,257 | 1,274 | 1,167 | 3,128 | 5,423 | 23,537 | 23,232 | 82,368 | 90,059 | 10,296 | 9,547 |

\*: TSD, target site duplications; IMD, insertion-mediated deletions

### SS-MEs impact genes in the primate genomes

To predict the functional impact of SS-MEs, we analyzed the gene context of their insertion sites based the gene annotation data in human from the GENCODE project (Release 23, July 2015) (Harrow, et al. 2012) combined with the NCBI RefGene annotation set (Pruitt, et al. 2007). Gene annotation information is lacking mostly for non-human genomes. In this analysis, for the non-human primate genomes, we used gene annotation based on the orthologous positions of human genes using the same liftOver data used in identifying the SS-MEs as described in the method section.

As shown in Table S7, a total of 46,466 SS-MEs, representing 20.1% of all SS-MEs, are located in genic regions, which include protein-coding genes, non-coding RNAs and transcribed pseudogenes. Similar to our observation for the human-specific MEs (Tang, et al. 2018), most of these genic SS-MEs (93.6%) are located in intron regions, while 1,926 SS-MEs contribute to exon regions as part of transcripts. Furthermore, these SS-MEs potentially impact the CDS regions of more than 90 unique genes, which cover all eight genomes (Table S8 & S9).

### DNA transposons were active during the later stage of primate evolution

DNA transposons, despite being quite active in the primate lineage until ∼37 Mya, are considered inactive in the later phase of the primate evolution (Pace Ii and Feschotte 2007). However, our data seem to suggest that DNA transposons still have a certain level of activity in all eight genomes analyzed in this study, which contributed a total of ∼2,400 (∼0.01% of all DNA transposons) DNA transposon entries being species-specific.

As shown in Table 2, SS-DNAs range from a few entries (8 in the crab-eating macaque genome) to a few hundred entries (more than 600 in the orangutan and gorilla genomes) in the eight primate genomes. Among these ∼2,400 SS-DNAs, 14.1% (340 entries) are also identified as the most recent SS-MEs (Table 2). Although not all these SS-DNAs are necessarily a result of canonical DNA transposon activity, they do provide strong evidence that some DNA transposons have remained active, albeit at a very low activity in the recent primate genomes.

## DISCUSSIONS

In this study, we deployed a comparative computational genomic approach recently developed for the analysis of human-specific MEs (Tang, et al. 2018) for a larger scale comparative genomic analysis involving eight primate genomes with four representing each of the top two families of primates, the *Hominoidea* and *Cercopithecoidea*. Our analysis provided the first set of comprehensive lists of MEs that are uniquely owned by each of these primate genomes based on the most updated reference sequences. Collectively, we identified a total of 230,855 SS-MEs from these eight primate genomes, among which 157,921 (68.4%) were considered to have occurred very recently in these genomes. These lists of SS-MEs and most recent SS-MEs allowed us to observe the differential DNA transposition and its impact in primate evolution. We discussed below the relevance of our results in several aspects.

### The challenges in the identification of SS-MEs

The reason for the lack of large-scale comparative studies for DNA transposition in primates is partly due to high challenges in this task. As previously discussed in our recent work about human-specific MEs (Tang, et al. 2018), identifying a comprehensive list of MEs uniquely owned by a primate genome faces certain challenges, which include but are limited to 1) the high content of MEs in the primate genomes, 2) the reference genome sequences are still incomplete, especially for the non-human primate genomes, and 3) genome assembly errors, especially for regions rich of repeat elements, which can mislead the results. For non-human primate genomes, we also face the lack of certain resources, for example, data linking the orthologous regions across closely related genomes (e.g. liftOver overchain files on the UCSC genome browser) and functional annotation data are mostly missing for comparative analysis among non-human primates. For these reasons, we believe that our lists of SS-MEs still suffer a certain level of false negatives and false positives. We can expect the situation to improve with continuing improvement of the genome assemblies, for example, benefiting from the use of newer generations of sequencing platforms that can provide much longer reads, such as the Nanopore and PacBio platforms (Roberts, et al. 2017; Schneider and Dekker 2012). The numbers of SS-MEs can be expected to have a certain level of increase from regions with sequencing gaps, especially regions highly rich of repeats, such as the centromere and telomere regions, which may be hot spots for certain types of MEs, such as LTRs (Tang, et al. 2018).

### The differential DNA transposition among primate genomes

Despite more and more non-human primate genomes have been sequenced and assembled in the recent years, prior studies on DNA transposition have mostly focused on the analysis of ME profiles for individual genomes separately (Battilana, et al. 2006; Ewing and Kazazian 2011; Jha, et al. 2009; Jordan, et al. 2018; Mills, et al. 2006; Ray, et al. 2005; Steely, et al. 2018; Stewart, et al. 2011; Tang, et al. 2018; Wang, et al. 2006). So far, only very limited comparative analyses involving a small number of genomes have been reported. Among these, the work by Mills et al (Mills, et al. 2006) compared the ME profile between human and chimpanzee, and a recent study has focused on lineage-specific *Alu* subfamilies in the baboon genome (Steely, et al. 2018). Due to the challenges described above, a large scale systematic comparative analysis of mobile elements in primate genomes still represents a gap in the field.

Our SS-ME data demonstrate that each primate genome display a remarkably different DNA transposition profile in terms of the overall amount of SS-MEs and the specific ME composition by ME class and subfamilies. Among the eight primate genomes examined, the number of SS-MEs in a genome varies from the highest at 66,578 copies in the baboon genome to the lowest at 3,281 copies in the crab-eating macaque genome (Table 2 and fig. 1). Overall, the *Hominidae* family has a lower level of SS-MEs 21,321 SS-ME/genome) than the *Cercopithecidae* family (36,393 SS-MEs/genome) (Table S5). While SINE represents the dominant class of SS-MEs for the *Cercopithecidae* family, L1s are more active in the *Hominidae* family, especially in the orangutan genome. SVAs as the youngest ME class are found to be uniquely in the *Hominidae* genomes, therefore verifying their unique association with *Hominidae* group proposed earlier without the genome sequences available (Wang, et al. 2005b) (Table 1). SVAs are shown to be the most active ME class in all *Hominidae* genomes and also among all ME classes for all eight genomes by the ratio of SS-MEs in the class, ranging from 16.6% in gorilla genome to 32.3% in orangutan genome, in comparison with no more than 2% for all other ME classes in any genomes (Table 3). Despite the presence of some sequences similar to SVAs in the *Cercopithecidae* genomes as MacSVAs, they are not active as shown by the numbers of SS-MacSVAs and most recent SS-MacSVAs from this class (Tables 2 and Table S4).

SS-MEs in each primate genome represent the total number of new MEs resulted from past DNA transposition since the divergence from the relative last common ancestor (LCA) among the species included in this analysis. Therefore, the number of SS-MEs in these primates can be impacted by the relative distance from their LCA, which are not the same among the eight primates. To avoid this bias, we also examined the most recent SS-MEs, which represents the number of most recent DNA transposition events in each genome independent of its evolution distance from other genomes, to compare the most recent level of recent DNA transposition across the genomes. As shown in Table 2 and fig. 1, despite the dramatic differences in the percentage of SS-MEs being most recent SS-MEs among the genomes, the overall pattern of relative DNA transposition across the genomes is still quite similar to that of SS-MEs. In the meantime, more detailed differences among the genomes were also revealed.

The crab-eating macaque genome has strikingly low numbers of both SS-MEs and most recent SS-MEs being ∼1/10 of that for averages across all eight primate genomes and ∼1/15 of *Cercopithecidae* family average (Table 2). This indicates that the extremely low level of DNA transposition in this genome was not due to a bias related to evolution distance, but a truly steadily extremely low DNA transposition. It is also interesting to note that, unlike other genomes, which have one or more classes of SS-MEs or most recent SS-MEs ranking high among the genomes, the crab-eating macaque genome has the least number of SS-MEs for all ME classes (fig. 1B & Table 2). This strongly suggests the existence of a molecular mechanism in this genome, which imposes a strong genome-wide suppression of DNA transposition. One possible such mechanism may be related to epigenetic regulation, such as a genome-wide DNA hyper-methylation during gametogenesis, as DNA methylation has been known to suppress DNA transposition (Law and Jacobsen 2010).

In contrast to the crab-eating macaque genome, the baboon genome has the largest number of SS-MEs and most recent SS-MEs that are more than 3 times higher than the genome averages with the majority contributed by SINEs (Table 2 & S5, fig. 1B & C). The number of SS-SINEs and most recent SS-SINEs in this genome is more than 4 times higher than the genome average (Table S5), reflecting SINEs being extremely successful in this genome. This confirms the previous observation that SINEs are quite active in the baboon genome (Steely, et al. 2018). By the average numbers of both SS-MEs and most recent SS-MEs, SINEs are more successful in the *Cercopithecidae* group than in the *Hominidae* group, while LINEs showed an opposite trend (Table S5).

The comparison of between the profile of SS-MEs and most recent SS-MEs across close-related genomes provides us with more details about the differences of DNA transposition among genomes. For example, between human and chimpanzee genomes, even though the latter has a higher number of SS-MEs for all ME types, the human genome has a much higher percentage of SS-MEs being the most recent SS-MEs (fig. 3). The largest difference is seen for SS-SINEs; while the human genome has significantly less SS-SINEs than the chimpanzee genome (8,844 vs. 10,612), it has more than double of the most recent SS-SINEs than in the chimpanzee genome (8,131 vs. 3,587) (Table 2). The percentage of the most recent SS-SINEs among SS-SINEs is >90% in the human genome compared to ∼30% in the chimpanzee genome (fig. 3B & C). This may suggest that, relatively speaking between the two genomes, DNA transposition was relatively lower in the human genome during the early stage; but accelerated more due to the emergence of a few very young and active SINE subfamilies, such as AluYa5, AluYb8, and AluYb9, along with L1HS, and SVA_D (fig. 5). These young and highly active ME subfamilies contributed to the higher percentage of most recent SS-LINEs and SS-SVAs in the human genome. It is worth noting that our lists of SS-MEs for human and chimpanzee (14,947 and 22,087, respectively) are not only significantly larger than the number of species-specific MEs reported in an earlier comparative study involving just a pairwise comparison between the same two genomes with earlier versions of the genome sequences (Mills, et al. 2006) (7,786 and 2,933, respectively), but also showing a different trend with the chimpanzee showing a larger number of SS-MEs in our study. This demonstrates the significant impact of the genome sequence quality, perhaps also the methodologies, on the results.

Similar to human genome, the baboon genome also has a very high recent activity of SINEs due to highly active subfamilies, such as AluRd4 and AluRd2 (fig. 5). The relative level of most recent DNA transposition measured based on the number of most recent SS-MEs in the eight primate genomes is clearly correlated with the numbers of the highly active ME subfamilies and their activity levels (fig. 5). Interestingly, in the human genome, the activity of the top five active ME subfamilies are all higher than any active ME subfamilies in the other genomes, revealing human as the youngest and most actively evolving species by its most active DNA transposition for the most recent period. All of the most active ME subfamilies below to the non-LTR retrotransposons which are all driven by the L1-based TPRT mechanism (Goodier 2016). This agrees with our recent observation that human genome has the largest number of functional L1s among primates and with most of these L1s being human-specific and even polymorphic (Nanayakkara, et al, manuscript in preparation). We believe that the largest number of functional L1s uniquely present in the human genome has provided the unique opportunities for the emergence of many young and active non-LTR ME subfamilies during human evolution.

In summary, our data indicate that the overall DNA transposition level among the eight primate genomes came as a result of differential activity levels of different ME classes and subfamilies and a different trajectories of activity level since the divergence from their perspective LCA.

### The impact of differential DNA transposition on primate genomes

The 230,855 SS-MEs from the eight primate genomes have collectively contributed to ∼82 Mbp net increase in the primate genomes (Table 4) with a net increase in each genome, ranging from ∼1.2Mbp in the crab-eating macaque genome to ∼25.5 Mbp in the orangutan genome. These amounts of genomic sequences are close to half of the human chromosome Y or 21 and is larger than the genomes of many free-living eukaryotic organisms, making DNA transposition as a very important, likely the most significant molecular mechansm contributing to genome size increases in primate genomes as previously discussed (Tang, et al. 2018).

In assessing the functional impact of these SS-MEs on genes, we had to address the lack of functional annotation for most of the non-human primate genomes. By using the human genome, the one with best functional annotation, as the reference and by identifying the orthologous regions of human genes in other genomes based on the same orthologous relationship data used in identification of SS-MEs, we were able to provide a prelimiary assessment of SS-MEs’s potential impact on genes in all of the non-human primate genomes.

Out results showed that a total of 46,466 SS-MEs, representing 20.1% of all SS-MEs, are located in genic regions, which include protein coding genes, non-coding RNAs, and transcribed pseudogenes (Table S7). This ratio is lower than the 50.7% previously reported for the human-specific MEs (Tang, et al. 2018), likely due to the lack of accurate gene annotation for the non-human primate genomes. Among these SS-MEs, 1,926 can potentially be part of the primate transcriptomes. Interestingly, in 92 of these cases, an SS-ME contributes to the protein coding sequence (CDS) in a transcript. Most of these CDS SS-MEs are SS-SINEs (40/92), even more so in the the *Cercopithecidae* family (33/38).

In summary, our data suggest that, similar to human-specific MEs (Tang, et al. 2018), SS-MEs in the primate genomes have the potential to participate in gene function via regulation of transcription, splicing, and contribution to protein-coding in a species-specific fashion. Certainly, the approach used here misses genes and transcript isoforms that are either species- or lineage-specific and not seen in the human genome. Therefore, we believe that our data represent an underestimation of SS-MEs’ impact on gene function in these genomes.

### Are DNA transposons still active in primate genomes?

DNA transposons have contributed to ∼3.6% of the primate genomes, yet their role in the primate evolutionary and their evolution profiles have yet to be extensively studied and fully understood. An early comprehensive study on DNA transposon activity concluded that DNA transposons, despite being very active during the early stages of primate evolution, have become inactive in the human genome after ∼37 Mya (Pace Ii and Feschotte 2007). However, this study was limited by the lack of diverse mammalian and primate genomic sequences at the time and the limited coverage to 794 DNA transposons nested within primate-specific *L1*/*Alu* elements.

Our comparative primate genome analysis identified a total of ∼2,400 DNA transposons as SS-MEs, representing ∼0.1% of all DNA transposons in these genomes with the numbers of SS-DNAs ranging from a few entries (8 in the crab-eating macaque genome) to a few hundred entries (668 in the orangutan genome) (Table 2). Among these SS-DNAs, 340 (14.1%) represent most recent SS-MEs, with the activity level for certain subfamilies, such as *MER97b*, being close to those of *ERVK* in certain genomes (Table S6 and fig. S3). While the existence of these SS-DNAs may be a collective result of the authentic cut-and-paste mechanism and several non-DNA transposition mechanisms, such as genomic rearrangement, transposition-based transduction, it might also suggest that DNA transposons are likely still active in these primate genomes, albeit at an extremely low activity.

### Conclusions and future perspectives

In summary, our comparative genomic analysis of eight primate genomes involving representatives from the top two primate families, *Hominidae* and *Cercopithecidae*, revealed remarkable differential levels of DNA transposition among primate genomes. Each of these genomes was shown to have a unique profile of SS-MEs in terms of their composition by ME class and activity level, and there are also common trends characteristic of lineages. Notably, the DNA transposition seems to be lowered to a ground level for all ME classes in the crab-eating macaque genome, likely due to a genome-wide suppression of DNA transposition, while it is highly active in the baboon and human genomes, each due to the existence of several unique highly active ME subfamilies. Overall, *Hominidae* has relatively more successful LINEs, while *Cercopithecidae* has SINEs as more successful. Remarkable differences in DNA transposition are also seen closely related genomes, as seen between human and chimpanzee genomes, with DNA transposition showing a later and quicker acceleration in the human genome compared to the chimpanzee genome. Furthermore, differential DNA transposition has made a significant differential impact on the genome size and gene function in these genomes, contributing to speciation and unique genomic and phenotypic characteristics of each species along with other mechanisms. Future studies may focus on elucidation of the specific mechanisms leading to such differential DNA transpositions in each species and the specific functional impacts on gene functions in context of the species-specific phenotypes.

## ACKNOWLEDGMENTS

This work is in part supported by grants from the Canadian Research Chair program, Canadian Foundation of Innovation, Ontario Ministry of Research and Innovation, Canadian Natural Science and Engineering Research Council (NSERC), and Brock University to PL, and was made possible by Compute Canada/SHARCNET high-performance computing facilities.

## A list of figures

**Fig. S1.**
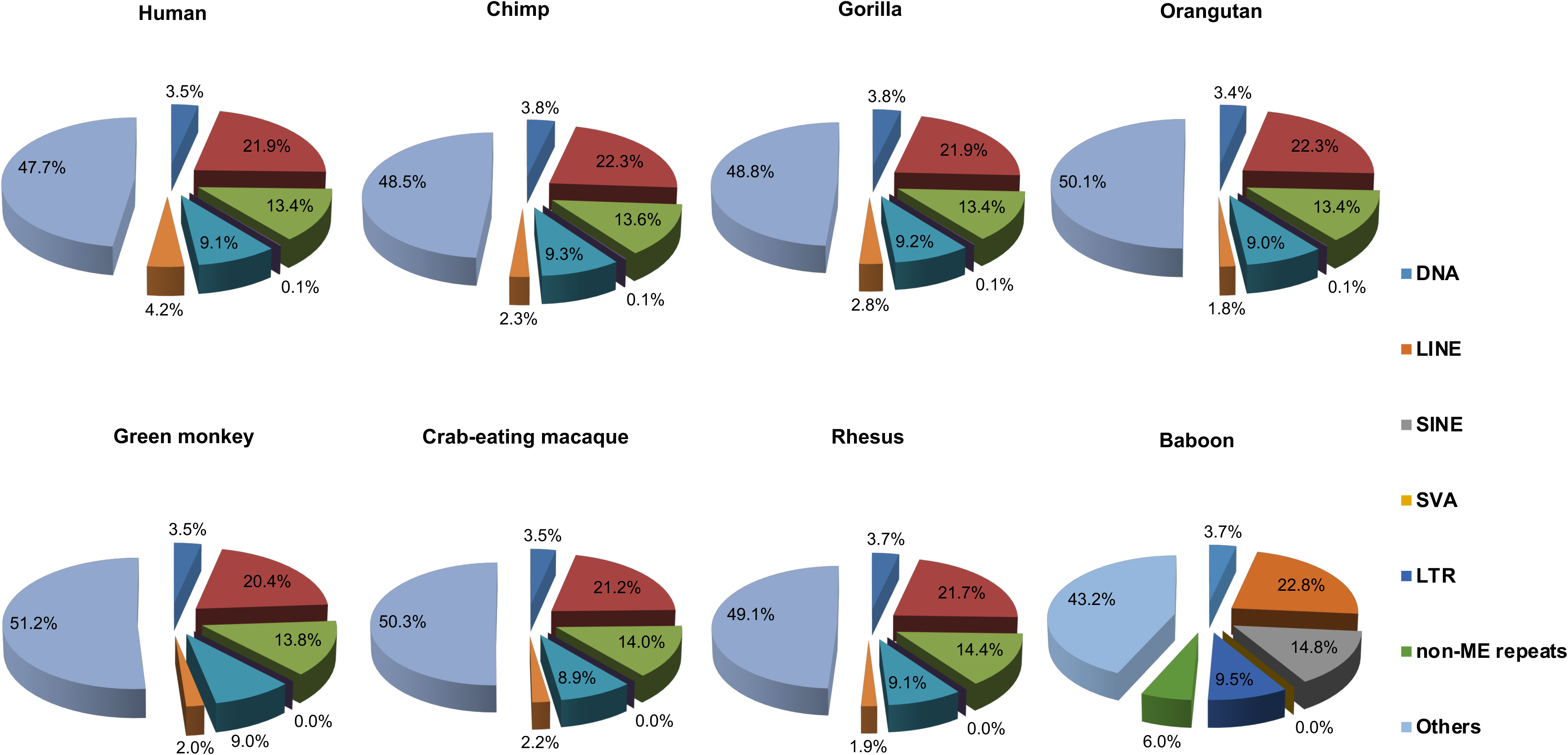
The repeat element composition by repeat class in eight primate genomes. Each pie chart shows the percentage of repeats by class in a genome. The ratios of genome components were based on sequence length. All gap regions in the eight primate genomes were excluded from the calculation.

**Fig. S2.**
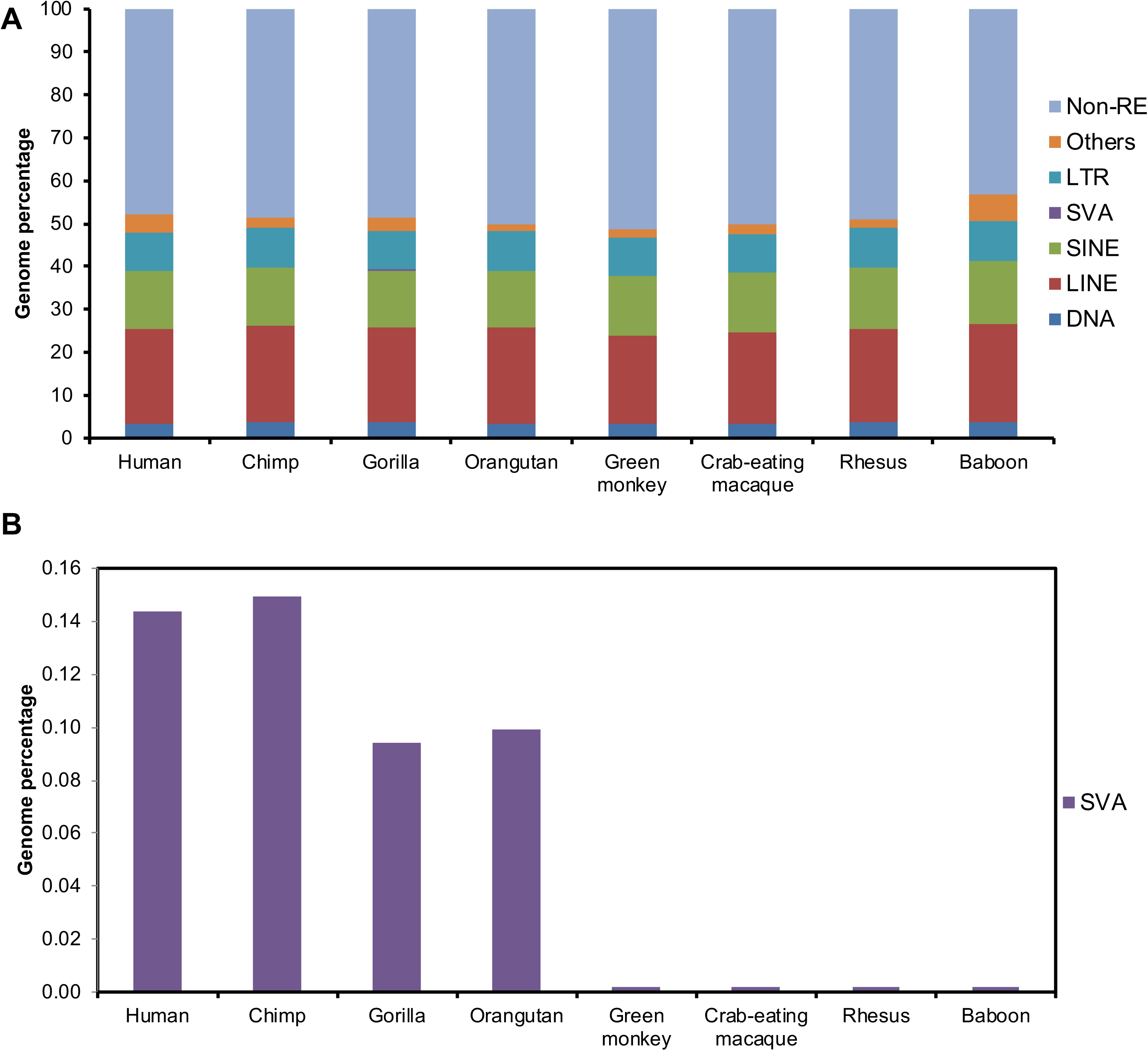
The composition of mobile elements (MEs) by ME class in eight primate genomes. Percentage of each ME class in a genome is calculated based on the size, total genome size is calculated non-gap sequences. A, the percentage of each ME class in genomes, including DNA, SINE, LINE, SVA, LTR, and Others in the primate genomes; B, Zoomed in section for the percentage of SVAs in in the genomes.

**Fig. S3.**
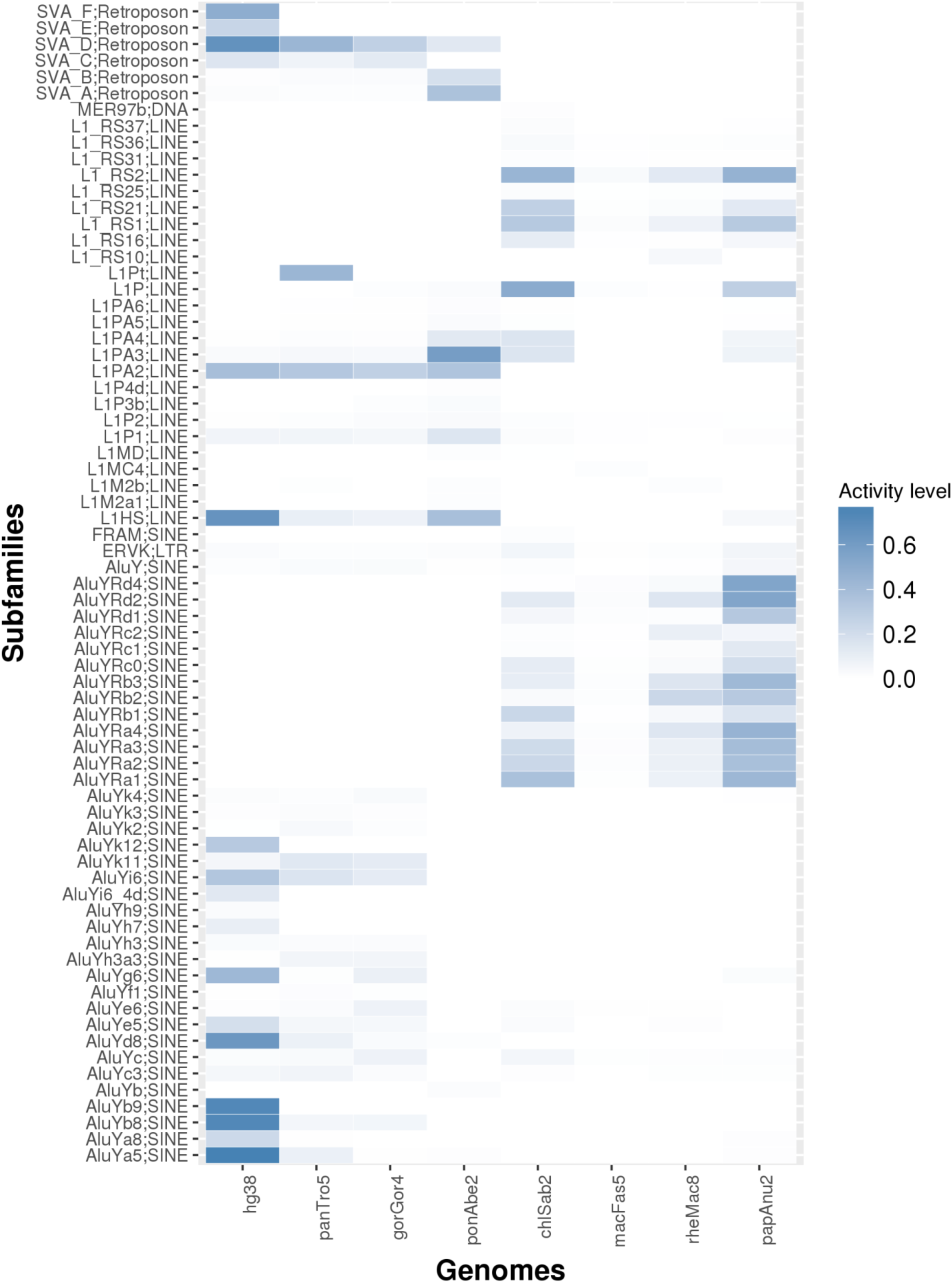
A heat map of mobile element (ME) subfamily activity in primate genomes based on most recent species-specific MEs. A total of 71 different ME subfamilies, which have activity (ratio of SS-MEs among all MEs in the subfamily) ≥1% in at least one genome, are selected and represented in the heat map. The 8 primate genomes are human(hg38), chimpanzee (panTro5), gorilla (gorGor4), orangutan (ponAbe2), green monkey (chlSab2), crab-eating macaque (macFas5), rhesus(rheMac8), and baboon (papAnu2). The activity level is calculated as the percentage of the most recent SS-MEs among the total number of MEs in the same subfamily in the genome. The detailed numeric values used to generate this heat map can be found in Table S6.

## A list of supplementary tables

**Table S1. Mobile element (ME) composition by sequence size in eight primate genomes. Table S2. Non-mobile element repeat profiles in eight primate genomes.**

**Table S2. Full-length mobile elements (MEs) before and after consolidation in eight primate genomes. Table S4. The copy numbers and lengths of SVA subfamilies in eight primate genomes.**

**Table S5. Average numbers of species-specific mobile elelements (SS-MEs) and most recent SS-MEs. Table S6. The most active mobile element (ME) subfamilies in eight primate genomes.**

**Table S7. The distribution of species-specific mobile elmements (SS-MEs) in the genic regions in eight primate genomes.**

**Table S8. Number of species-specific mobile elements (SS-MEs) in potential protein coding genes in eight primate genomes.**

**Table S9. A list of species-specific mobile elements (SS-MEs) in potential protein coding genes in eight primate genomes.**

